# waveRAPID® - a robust assay for high-throughput kinetic screens with the Creoptix® WAVEsystem

**DOI:** 10.1101/2021.02.05.429874

**Authors:** Önder Kartal, Fabio Andres, May Poh Lai, Rony Nehme, Kaspar Cottier

## Abstract

Surface-based biophysical methods for measuring binding kinetics of molecular interactions, such as Surface Plasmon Resonance (SPR) or Grating-Coupled Interferometry (GCI), are now well established and widely used in drug discovery. Increasing throughput is an often-cited need in the drug discovery process, and this has been achieved with new instrument generations where multiple interactions are measured in parallel, shortening the total measurement times and enabling new application areas within the field. Here, we present the development of a novel technology called waveRAPID for a further - up to ten-fold - increase in throughput, consisting of an injection method using a single sample. Instead of sequentially injecting increasing analyte concentrations for constant durations, the analyte is injected at a single concentration in short pulses of increasing durations. A major advantage of the new method is its ability to determine kinetics from a single well of a micro-titer plate, making it uniquely suitable for kinetic screening. We present the fundamentals of this approach using a small molecule model system for experimental validation and comparing kinetic parameters to traditional methods. By varying experimental conditions, we furthermore assess the robustness of this new technique.

Finally, we discuss its potential for improving hit quality and shortening cycle times in the areas of fragment screening, low molecule weight compound screening, and hit-to-lead optimization.

## Introduction

The accurate and efficient measurement of intermolecular interactions and binding events is a critical element for all manner of basic research and is indispensable for drug discovery programs. Ligand binding assays can be performed using labeled molecules (radiolabels, fluorescent labels, etc.), but non-disruptive labeling and elaborate washing and purification steps are often required. The kinetics of interactions are also not easily acquired or understood using these methods. Refractometric techniques such as Surface Plasmon Resonance^1^ (SPR), in contrast, allow the monitoring of interactions in real time using unlabeled species. However, SPR measurements usually require significant time investments, as one interaction partner needs to be covalently bound near the sensor surface while the other must be injected repeatedly at different concentrations. SPR sensors are also inherently sensitive to bulk refractive index changes. These issues limit the utility of SPR in screening programs. Surface-based optical detection of molecular interactions has evolved in the past years, and with the introduction of Grating-Coupled Interferometry^2–4^ (GCI) some typical limitations of SPR, such as sensitivity and sample robustness, have been mitigated.

In a typical experiment for measuring kinetics, the first interaction partner (the so-called ligand) such as a drug target is immobilized or captured on the sensor surface. Then, the second interaction partner (the analyte) is repeatedly injected at increasing steady concentrations, each of which is applied for the same duration. Both SPR and GCI measure small changes in the refractive index near the sensor. When an analyte binds, the refractive index changes proportionally to the mass of the interacting analyte. After each injection, the signal is often allowed to return to baseline in the absence of analyte (sometimes forcibly with regeneration conditions that disrupt binding). Alternatively, the analyte concentration can be increased steadily in a regeneration-free experiment. Binding affinity (*K*_*d*_) and kinetic parameters (*k*_*a*_, *k*_*d*_) are obtained from the resulting response signal (Figure 1, top).

**Figure 1.**
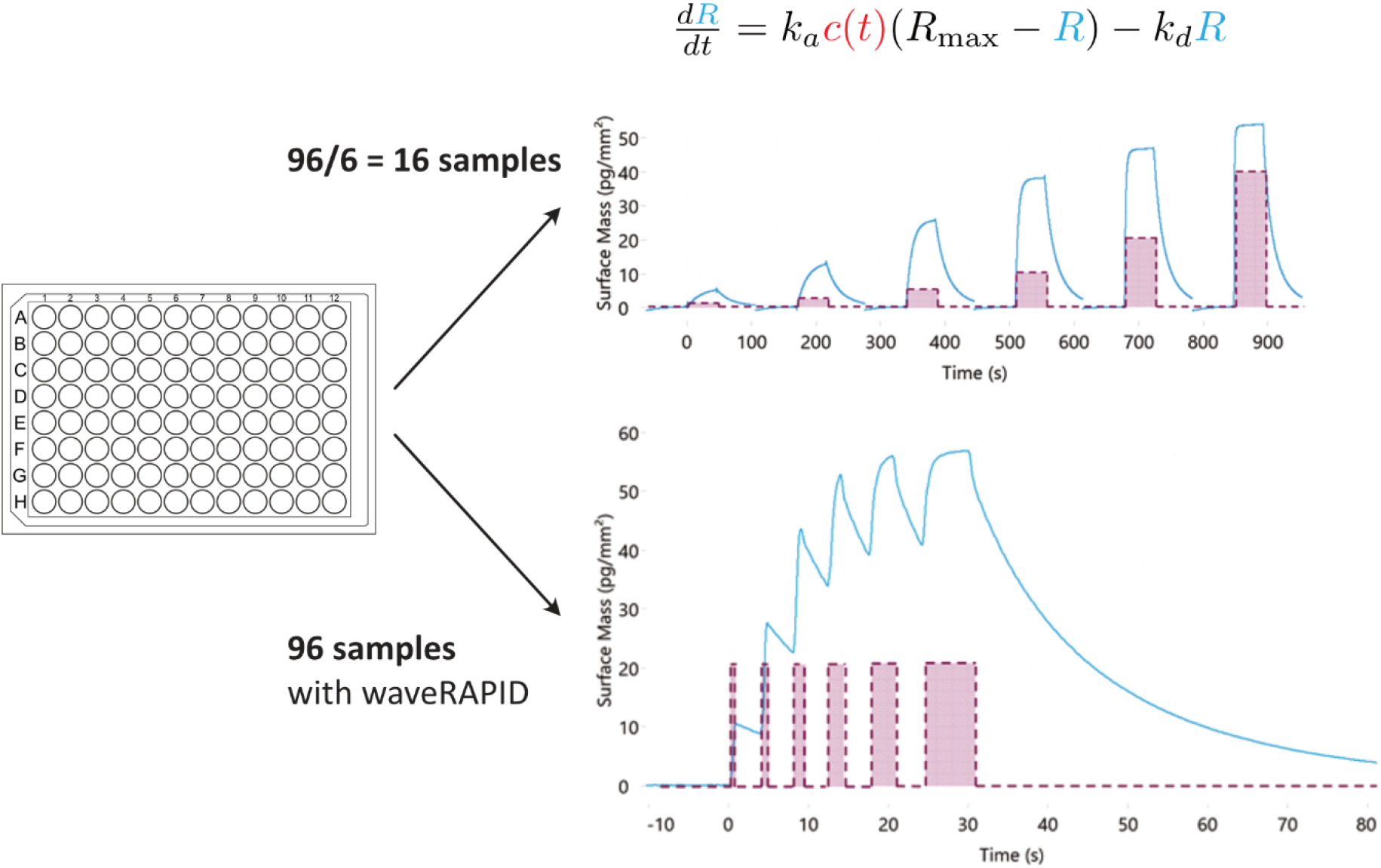
Traditional vs waveRAPID kinetics. In a typical ligand binding assay (top), the analyte is introduced at increasing concentrations, with each injection being of uniform duration. In each cycle, the injected concentration *c*_*i*_ is fixed but *c*(*t*) is a function of time for the whole series (red). *R* is the the sensorgram response, proportional to the mass of bound analyte (blue). Model fitting provides estimates for the parameters (*k*_*a*_, *k*_*d*_, and *R*_*max*_) of the ODE for one-to-one binding (top) that best explain the data in all segments of the time series. In the waveRAPID assay, the series consists of a single cycle with only one association phase that is interrupted by dissociation portions and a final dissociation phase. Model fitting can be done with the dissociation segments only.

A crucial step in the drug discovery process is the screening of many compounds to narrow down the candidates to a few promising leads^5^. This requires rapid characterization of binding interactions while avoiding false positives that slow down the discovery process considerably. Traditional kinetic assays with multiple concentrations (multi-cycle) are time intensive and therefore not suited for screening thousands of candidate compounds. Measurements with a single injection at a single concentration (single cycle) are faster but do not yield reliable estimates of all kinetic parameters.^6–9^ One way to increase throughput is to run several traditional, multi-cycle assays in parallel. This has been achieved with new generations of instruments that shorten the total measurement time and thereby enable new applications of label-free kinetics. However, brute-force parallelization leads to an explosion of reagent and buffer consumption and therefore to an increase in running costs that may become prohibitive when screening many compounds.

In a different, more elegant approach, Shank-Retzlaff and Sligar^10^ as well as Quinn^11,12^ have shown that single injections producing a time-dependent concentration gradient of the analyte at the sensor surface can in principle replace multi-cycle assays without serious detrimental effects on parameter estimation. However, Shank-Retzlaff and Sligar noted a marked sensitivity of their assay to instrument noise. Quinn’s method, which has been commercialized in the ForteBio Pioneer Systems as OneStep injection technology, relies on natural diffusion and is therefore sensitive to variations of the analyte’s diffusion constant. Furthermore, the sensorgrams of both assays are difficult to interpret, which has limited acceptance in the market.

Here, we introduce a new alternative kinetic assay^13^, Repeated Analyte Pulses of Increasing Duration (waveRAPID), which – similarly to the above-mentioned methods – relies on a time-dependent concentration profile of the analyte at the sensor surface. However, in contrast to the monotonously increasing concentration profiles found in previous methods, waveRAPID generates a pulsating concentration profile by injecting the analyte at the same concentration but multiple times. The reliable characterization of an interaction using a single pickup from a single well, avoiding the typical washing steps between injections, is a significant boost for screening applications. For example, 96 different interactions can be analyzed from a single 96-well microtiter plate (Figure 1) as compared to only 16 with a traditional assay using a six-fold dilution series.

## Materials and Methods

### Implementation of the waveRAPID assay

The waveRAPID assay has been designed to achieve high-quality estimates of kinetic parameters with less preparation and measurement time by using the same concepts and hardware as traditional label-free kinetic measurements. To achieve an informative binding signal, the WAVEsystem applies multiple short pulses of the analyte at a single concentration but of increasing durations. In contrast to traditional kinetics, where a single cycle involves injection of the analyte followed by a single dissociation phase (Figure 1, top), a single cycle of waveRAPID combines multiple association phases with multiple intermittent dissociation segments, concluded by a longer dissociation phase (Figure 1, bottom). This unique injection pattern drives the binding response to extract as much information as possible from a single concentration input.

To achieve the concentration function *c*(*t*) driving the waveRAPID assay, the microfluidic system must allow fast transitions between analyte and buffer flows. The microfluidics of the Creoptix WAVEsystem is very well suited for waveRAPID assays. Disposable cartridges with parallel flow channels and valves enable ultra-fast transition times of 150 ms and prevent clogs. The no-clog microfluidics makes the direct study of biofluids and crude reaction mixtures possible, and the system is compatible with or tolerant to harsh, uncommon solvents such as acetonitrile or high concentrations of DMSO. Thus, the Creoptix WAVEsystem supports the application of the RAPID technique in a variety of use cases.

Despite the capabilities of the WAVEsystem, physical constraints rule out ideal unit pulses as concentration functions. In fact, the driving force *c*(*t*) for the binding reaction is not the injected concentration but the concentration at the sensor surface which is governed by convection and diffusion^14,15^, leading to a flowrate-dependent dispersion characteristics of the fluidics. Assuming a simple one-to-one kinetic model, an ideal timing for the pulse durations can be determined by considering a) injected concentration, b) the dispersion characteristics of the fluidics, c) the acquisition rate, as well as d) the lowest possible binding levels that can be measured in association, and finally e) the limitations for measuring slow dissociations due to drift. These theoretical considerations, as well as simulations and experimental validation, suggest a sequence of six pulses with durations that are defined *relative* to the total injection duration (the duration from the start of the first pulse to the end of the last pulse). The software computes the *absolute* pulse durations by multiplying the relative durations by the user-specified total injection time. For user convenience, three parameter presets are available in WAVEcontrol based on the range of affinity of interest. Table 1 lists the parameters of these presets, including flowrate, acquisition rate, analyte concentration, and pulse timing. With these presets, we try to ensure that mass-transport is not limiting, which occurs particularly at high immobilization levels in combination with fast association rates and low flow rates. Nonetheless, as Shank-Retzlaff and Sligar^10^ have shown, the binding signal driven by a time-dependent concentration function *c*(*t*) can be analyzed using a two-compartment model for taking these effects into consideration. This functionality is also available in WAVEcontrol.

**Table 1.** Pulse durations for different presets in WAVEcontrol.

|  | Weak Binders<br>(1mM-100mM) | Intermediate<br>Binders<br>(5nM-5mM) | Tight Binders<br>(1pM-20nM) |
| --- | --- | --- | --- |
| Flowrate | 400 mL/min | 100 mL/min | 50 mL/min |
| Acquisition Rate | 40 Hz | 10 Hz | 1 Hz |
| Analyte<br>Concentration | 200 mM | 10 mM | 500 nM |
| 1<sup>st</sup> Association duration | 0.031 s | 0.156 s | 0.750 s |
| 1<sup>st</sup> Dissociation Duration | 0.606 s | 3.031s | 14.550 s |
| 2<sup>nd</sup> Association Duration | 0.063 s | 0.313 s | 1.550 s |
| 2<sup>nd</sup> Dissociation Duration | 0.606 s | 3.031 s | 14.550 s |
| 3<sup>rd</sup> Association duration | 0.125 s | 0.625 s | 3.000 s |
| 3<sup>rd</sup> Dissociation Duration | 0.606 s | 3.031 s | 14.550 s |
| 4<sup>th</sup> Association Duration | 0.250 s | 1.250 s | 6.000 s |
| 4<sup>th</sup> Dissociation Duration | 0.606 s | 3.031 s | 14.550 s |
| 5<sup>th</sup> Association Duration | 0.500 s | 2.500 s | 12.000 s |
| 5<sup>th</sup> Dissociation Duration | 0.606 s | 3.031 s | 14.550 s |
| 6<sup>th</sup> Association Duration | 1 s | 5 s | 24 s |
| Last Dissociation Duration | 20 s | 300 s | 1800 s |
| Total Duration | 25 s | 325 s | 1920 s |

For the model-based data evaluation, the time-dependent concentration profile near the sensor surface must be known. Conveniently, this function can be measured directly by GCI during a DMSO calibration step, which consists in a waveRAPID cycle using the same parameters as the actual measurement. Here, in the absence of binding, GCI is effectively used as a concentration sensor. Since the evanescent wave extends up to the order of 100-200 nm into the bulk solution^16^, only the concentration in the unstirred layer of the laminar flow close to the sensor surface is measured. This reference concentration measurement is representative of the concentration of an analyte compound in the absence of binding if the diffusion properties are similar. To obtain the function *c*(*t*) used in the fitted model, the reference concentration profile is normalized (i.e., divided by the maximum reference response) and scaled (i.e., multiplied by the input concentration of the analyte used in the measurement).

### Parameter estimation using Direct Kinetics

Data analysis was performed using WAVEcontrol (version 4.0). Reference channel subtraction was performed to correct for bulk effect and non-specific binding. In WAVEcontrol 4.0, the kinetic parameters are estimated with our new, patented Direct Kinetics engine^17^ which allows complete data analysis with a single mouse click. Direct Kinetics uses an intermediate basis function model for the sensorgram time series to obtain an approximating interpolation. The fitted interpolant can be used to obtain initial parameter estimates^18^ of the underlying ordinary differential equation (ODE). The estimates are then further improved using generalized profiling^19^. The ODEs used to evaluate waveRAPID data are the same as in traditional kinetics but the time-dependent input concentration acts as a driving force that makes the ODEs non-autonomous.

Refractive index (RI) disturbances are avoided by setting the weights of the data residuals in the association parts to zero (setting “Point weights” in WAVEcontrol 4.0). That is, the misfit in the association parts of the pulses does not impact the parameter estimation. Nevertheless, the starting point of each pulse decay depends on the previous increase of the binding signal. Therefore, the dissociation curves are determined by and must retain the information about *k*_*a*_ and *R*_*max*_. Experiments have shown that using a set of subsequent dissociation parts that result from different analyte injection times provides enough information to fit the complete model and reliably estimate all parameters, including *k*_*a*_ and *R*_*max*_.

### FUR binding to CAII on 4PCH WAVEchip^®^

Carbonic anhydrase (CAII) was immobilized onto the 4PCH sensor chip surface by the amine coupling method. Briefly, the polycarboxylate hydrogel layer on the 4PCHP sensor chip surface was activated with EDC/NHS solution. CAII samples (0 µg/mL, 5 µg/mL, 10 µg/mL, and 25 µg/mL) were prepared in sodium acetate (pH 5) and injected to the surface at a flow rate of 10 µL/min. The final surface density for each non-reference channel was 600 pg/mm^2^, 2000 pg/mm^2^, and 14000 pg/mm^2^, respectively. After the immobilization, the sensor chip surface was blocked using 1M ethanolamine hydrochloride (pH 8.0).

For assessing the reproducibility of the waveRAPID assays, three concentrations (5.5 µM, 17µM, and 50 µM) of furosemide (FUR) were prepared using assay buffer (PBS containing 3% DMSO). The experiments were carried out at flow rate of 80 µL/min. Briefly, 120 µL of FUR samples were injected for 20 s or 30 s total injection duration, followed by a 60 s dissociation with assay buffer.

For the traditional kinetic assay, FUR was serially diluted 1:3 using assay buffer for a six-point dose curve (50 µM, 17 µM, 5.5 µM, 1.8 µM, 0.6 µM, 0.2 µM). The experiment was carried out at flow rate of 80 µL/min. 100 µL of compounds were injected for 45 s, followed by a 60 s dissociation with assay buffer.

### Kinetic screening of compounds

Two undisclosed His-tagged target proteins (1 and 2) were immobilized on a 4PCP-NTA sensor chip surface using His capturing. Briefly, the NTA surface was charged with 0.5mM NiCl^2^ solution. The target protein 1 (10 µg/mL) and target protein 2 (50 µg/mL) were prepared in PBS and injected to the surface at flow rate of 10 µL/min. The final surface density was 4000 – 5000 pg/mm^2^. A set of 90 compounds (at 10 µM or 50 µM) were prepared using assay buffer (50 mM Phosphate Buffer, 0.05% Tween, 5% DMSO). The compounds were injected for 25 s, followed by 250 s dissociation with assay buffer.

## Results

### waveRAPID data enables estimation without RI artifacts

The waveRAPID data can be evaluated with any appropriate dynamic model, for example the ODE illustrated in Figure 1 which applies to a simple one-to-one interaction. Conveniently, the concentration function *c*(*t*) can be extracted from the calibration step with a reference molecule such as DSMO which requires only a single injection, even if many compounds are measured. The design of the assay also entails an improvement in parameter estimation by avoiding the data affected by refractive index (RI) disturbances.

When fitting a model to waveRAPID data, it is sufficient to minimize only the misfit in the dissociation part of each pulse when no analyte should be present in the bulk phase. This ability overcomes a fundamental drawback of refractive index sensors where the presence of unbound analyte in the buffer can significantly alter the binding response signal.

### waveRAPID estimates are consistent and reproducible

To demonstrate the robustness and reproducibility of the waveRAPID method, we performed a kinetic analysis of the small molecular analyte furosemide, binding to Carbonic Anhydrase II (CAII) over a range of different conditions. Figure 2 compares a single waveRAPID measurement against traditional multi-cycle kinetic measurements, confirming that both methods deliver highly consistent results. To confirm that waveRAPID estimates are reproducible, we have repeated the same measurement at three different ligand densities, injecting the analyte at three different concentrations and for two different total injection durations (20 s and 30 s). All 18 measurements were completed within 20 minutes, and the double-referenced response data were fitted to a one-to-one binding model with WAVEcontrol’s Direct Kinetics engine^17^. The estimation of the kinetic parameters reveals a very good reproducibility (Figure 3) with relative standard errors at approximately 5% or below (Table 2). The estimated mean (sample standard deviation) of the affinity constant over all measurements is *K*_*d*_ = 1.71(0.16) µM.

**Table 2.** Kinetic parameter estimates for the interaction of furosemide with Carbonic Anhydrase II (CAII) measured at different conditions. The parameters are the maximum-likelihood estimates (MLEs) and the error is the standard error relative to the MLEs measured in %.

| Concentration (μM) | Time (s) | $k_a$ (M<sup>-1</sup>s<sup>-1</sup>) | $k_a$ error (%) | $k_d$ (s<sup>-1</sup>) | $k_d$ error (%) | $K_d$ (μM) | $R_{max}$ (pg/mm<sup>2</sup>) |
| --- | --- | --- | --- | --- | --- | --- | --- |
| 5 | 20 | 3.7E+04 | 6.0 | 5.6E-02 | 4.3 | 1.5 | 1.5 |
| 5 | 20 | 2.8E+04 | 7.3 | 5.2E-02 | 4.2 | 1.8 | 2.5 |
| 5 | 20 | 2.9E+04 | 2.4 | 5.6E-02 | 1.4 | 1.9 | 57.0 |
| 5 | 30 | 3.6E+04 | 4.2 | 5.2E-02 | 3.4 | 1.4 | 1.4 |
| 5 | 30 | 3.0E+04 | 4.0 | 5.2E-02 | 2.8 | 1.7 | 2.3 |
| 5 | 30 | 3.2E+04 | 1.9 | 5.5E-02 | 1.4 | 1.7 | 51.9 |
| 17 | 20 | 3.7E+04 | 4.2 | 5.6E-02 | 5.2 | 1.5 | 1.5 |
| 17 | 20 | 3.6E+04 | 5.8 | 5.1E-02 | 7.4 | 1.4 | 2.2 |
| 17 | 20 | 3.4E+04 | 1.5 | 5.9E-02 | 1.8 | 1.7 | 50.5 |
| 17 | 30 | 3.7E+04 | 3.3 | 5.8E-02 | 3.3 | 1.6 | 1.5 |
| 17 | 30 | 3.3E+04 | 3.8 | 5.3E-02 | 4.0 | 1.6 | 2.3 |
| 17 | 30 | 3.2E+04 | 1.5 | 5.8E-02 | 1.5 | 1.8 | 50.8 |
| 50 | 20 | 3.1E+04 | 2.8 | 5.5E-02 | 3.8 | 1.8 | 1.6 |
| 50 | 20 | 2.9E+04 | 3.4 | 5.2E-02 | 4.7 | 1.8 | 2.5 |
| 50 | 20 | 3.2E+04 | 1.5 | 5.9E-02 | 2.0 | 1.9 | 51.9 |
| 50 | 30 | 3.3E+04 | 4.6 | 5.7E-02 | 4.6 | 1.8 | 1.6 |
| 50 | 30 | 2.8E+04 | 5.0 | 5.4E-02 | 5.2 | 1.9 | 2.5 |
| 50 | 30 | 3.1E+04 | 2.1 | 5.8E-02 | 2.0 | 1.9 | 51.7 |

**Figure 2.**
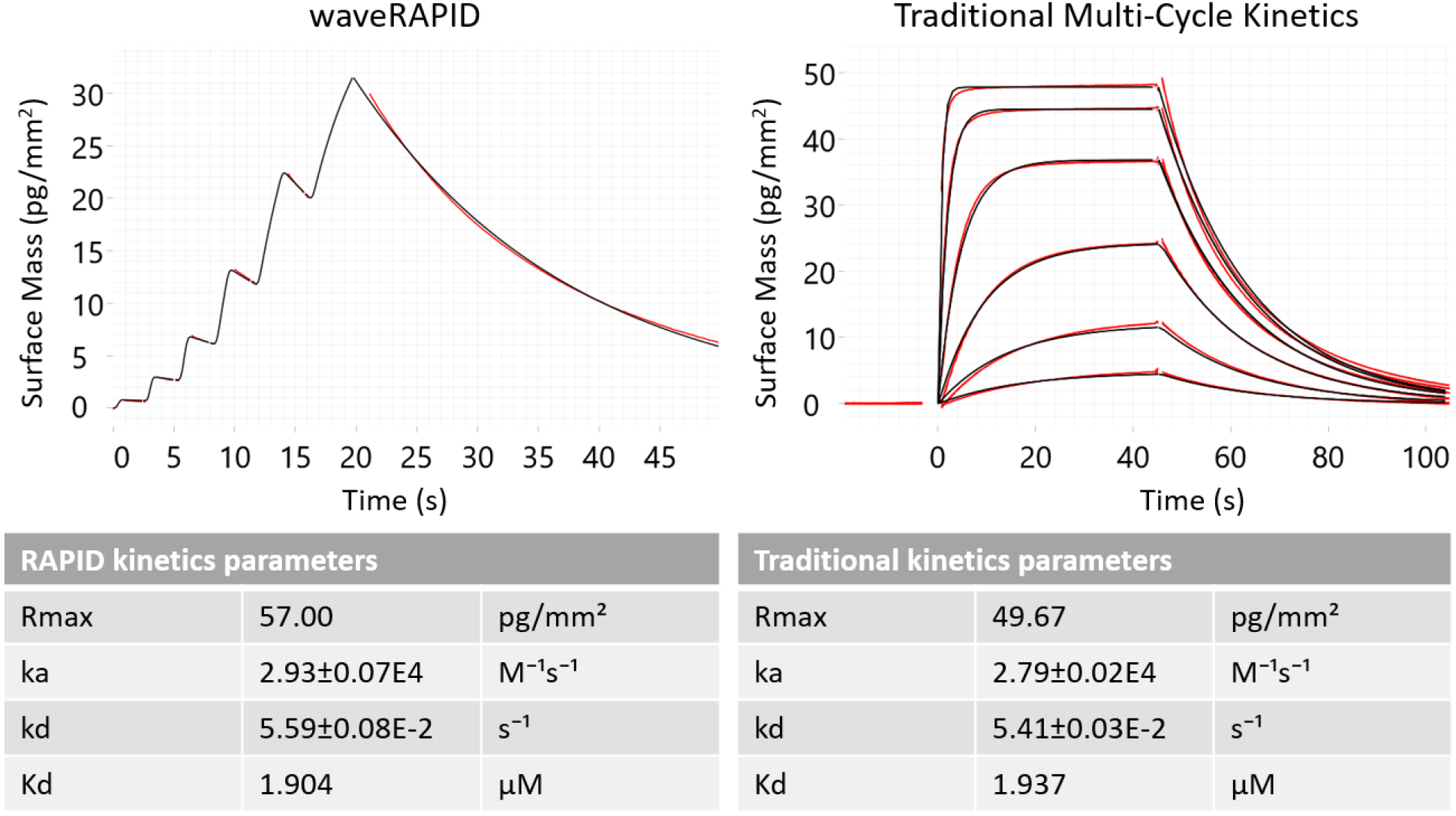
Comparison of traditional kinetic data and estimates with waveRAPID. Binding was measured for the small molecular compound furosemide (analyte) to Carbonic Anhydrase II (CAII, ligand) generated with either waveRAPID (left panel) or with traditional multi-cycle kinetics (right panel). The double-referenced response data (red) are fitted with a one-to-one binding model (black lines) using the Direct Kinetics engine of WAVEcontrol. Maximum-likelihood estimates (MLEs) of the kinetic parameters with 95% confidence limits are shown in the corresponding tables.

**Figure 3.**
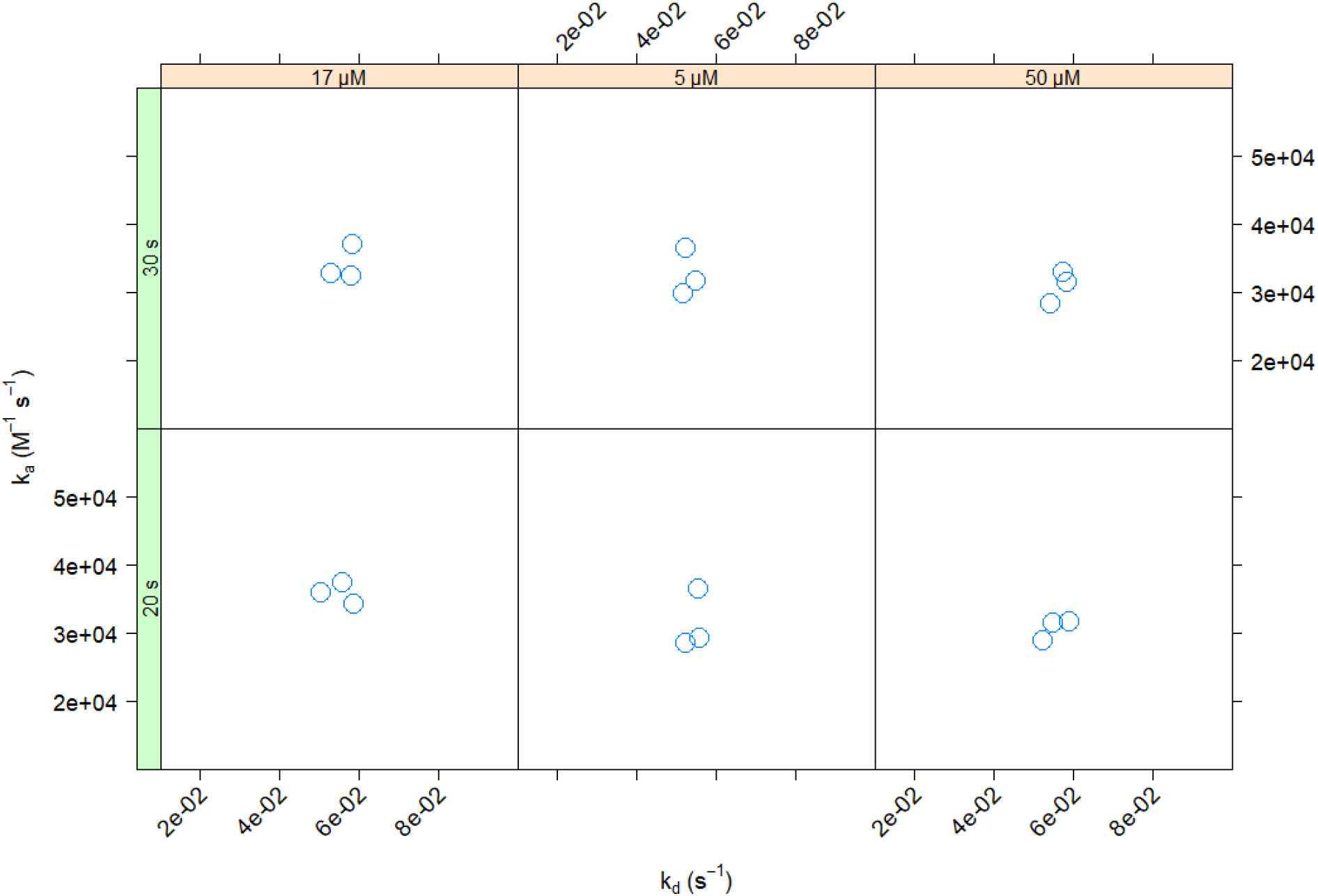
Reproducible estimates of kinetic constants for furosemide-CAII interaction. Each subfigure contains measurements at three different ligand densities for a certain combination of injection duration (left strip: 20 or 30 seconds) and analyte concentration (upper strip: 5, 17, or 50 μM).

### waveRAPID allows high throughput screening

The waveRAPID technology is suited for accelerating kinetic measurements, thus allowing high-throughput kinetic determination of many analytes in a screening setup. Because assays can be completed in significantly less time, inherently unstable species can more easily be studied. We performed a kinetic characterization of small molecule drug hits for an undisclosed drug target using 90 different analytes, each applied for 25 s of total injection duration prior to 300 s of dissociation. In just 18 hours of total assay time, several hits were successfully identified with affinity constants ranging from nM to µM concentrations (Figure 4). Compounds which showed no difficulties in previous assays as well as replicates of the control compound showed low estimation errors (data not shown). But with waveRAPID we could also evaluate compounds that proved to be difficult to analyze in traditional assays due to large bulk refractive index contribution. The remaining problematic compounds were clearly identified by large estimation errors. With its ability to detect weak binding interactions more reliably, the waveRAPID assay is optimally suited for small molecule compound screens and even fragment-based screens.

**Figure 4.**
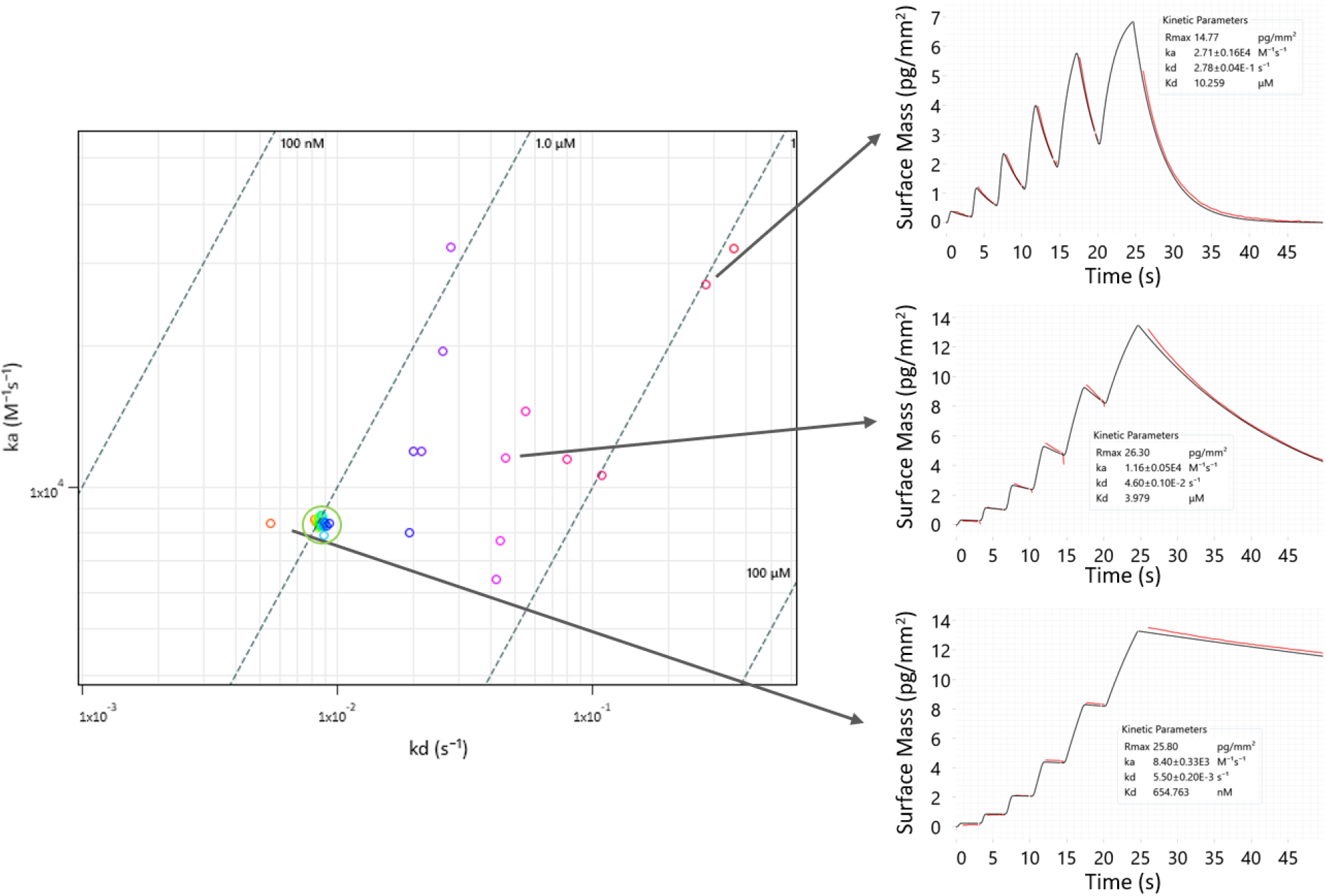
waveRAPID kinetic screen of small molecular drug hit candidates. Ninety molecules were analyzed, producing kinetic data in 18 hours. The results shown in the rate map were filtered for low statistical errors on the rate constants. Three interactions with different binding strengths are highlighted by showing sensorgrams and fits from WAVEcontrol.

## Discussion

We have shown that waveRAPID enables kinetic screening by redesigning the binding assay to significantly reduce the time for preparing as well as performing the measurements. Not only does this allow for higher throughput, but it also mitigates certain error sources: while using a single instead of multiple wells reduces the impact of pipetting mistakes, shorter experiments avoid measurement artifacts due to degradation of the sensor surface and unstable compounds. We argue that the low standard deviations we observed in our reproducibility tests could also partly be due to the simple reduction in total measurement time, reducing the lowest frequencies that are most impacted by 1/f noise^20^. To implement waveRAPID, we use existing hardware, in particular the capabilities of the WAVEsystem to measure rapidly changing signals with a high signal-to-noise ratio thanks to the GCI sensor. This combination of features is crucial to implement rapid kinetic screens that probe the interaction with short analyte pulses. Like gradient injections^10,11^, a single waveRAPID assay performs a full titration without the need for sensor regeneration and therefore provides informative sensorgrams that allow for reproducible estimates of the kinetic parameters including their uncertainties.

We have shown that the parameter estimates of waveRAPID assays are consistent with traditional assays (Figure 2) and robust towards different experimental configurations (Figure 3). These evaluations were done with the default one-to-one model, assuming no mass-transport limitation. However, the Direct Kinetics engine is capable to perform the evaluation with a two-compartment model as well. In general, the presets in WAVEcontrol (Table 1) try to avoid the effects of mass-transport limitation and for the cases that we have presented in this study, the one-to-one model seems appropriate. A more extensive theoretical and experimental comparison between the estimates of traditional and waveRAPID assays, wherein mass-transport effects are considered, is in preparation but outside the scope of this brief communication.

waveRAPID also makes it possible for the first time to evaluate a sensorgram without the parts that are affected by the RI of the bulk analyte. Hitherto, this was a major disadvantage of RI sensors, especially when evaluating slow reactions where a significant fraction of the analyte can be present in solution. The common solution to RI perturbations is to include this nuisance parameter in the fitted model as an additive effect. While this is possible, it is best practice to optimize the assay rather than complicating the model which always introduces additional assumptions that can break down in certain situations. A more complicated model tends to make it harder to reliably estimate the parameters because they become less identifiable^21^, especially if some input parameters, like the analyte concentration, are assumed to be constant^22,23^.

Arguably, the most important application of waveRAPID is in kinetic screening settings. In the kinetic screen that we have presented here as a proof-of-concept (Figure 4), the difficult compounds caused major workflow limitations in the traditional multi-cycle kinetics, making regular surface regeneration necessary, whereas waveRAPID enabled a seamless workflow with continuous unattended measuring time. The ranking based on the statistical error estimates confirmed the previously known, problematic compounds. Thus, hit quality is improved, as the statistical error estimates can be used to identify false positives that would otherwise slow down the discovery pipeline. Since waveRAPID delivers a full kinetic characterization of compounds with a single injection, kinetic screens now become possible. Thus, the information that in a classic screening setting would have been gathered in a secondary screen or hit confirmation step, is already available after one pass. Combining first and secondary screens into one single step has a real potential to change workflows in drug discovery.

In summary, the novel waveRAPID assay leverages traditional equipment and techniques but offers several compelling advantages due to a design that is superior to traditional assays. The increased throughput can improve hit quality and shorten cycle times considerably in early-stage small molecule drug discovery.

## Acknowledgments

We thank Idorsia Pharmaceuticals Ltd (Geoffroy Bourquin, Laksmei Goglia, Solange Meyer, and Oliver Peter) for providing the experimental data of the kinetic screen.

